# Intact maternal buffering of stress response in infant rats despite altered responsivity towards maternal olfactory cues in the valproic acid model of autism-like behavior

**DOI:** 10.1101/2022.03.27.485976

**Authors:** Amanda M. White, Xianli An, Jacek Debiec

**Author notes:** These authors contributed equally to this work (Amanda M. White, Xianli An). **Correspondence:** (AMW), (XA), (JD) – Corresponding Author. **Declaration of Interests:** None.

## Abstract

Disrupted processing of social cues and altered social behaviors are among the core symptoms of autism spectrum disorders (ASDs), and they emerge as early as the first year of life. These differences in sensory abilities may affect the ability of children with ASDs to securely attach to a caregiver and experience caregiver buffering of stress. Prenatal exposure to valproic acid (VPA) has been used to model some aspects of ASDs in rodents. Here, we asked whether prenatal VPA exposure altered infant rats’ behavioral responsivity to maternal olfactory cues in an odor preference test and affected maternal buffering of infants’ stress responsivity to shock. In the odor preference test, one-week old rats treated with VPA during pregnancy appeared to have impaired social recognition and/or may be less motivated to approach social odors in early infancy. These effects were particularly prominent in female pups. In two-week old rats, VPA-exposed pups and saline-exposed pups showed similar preferences for home cage bedding. Although VPA-exposed pups may initially have a deficit in this attachment-related behavior they do recover typical responses to home cage bedding in later infancy. Both control and VPA-exposed pups showed robust stress hormone responses to repeated shocks, an effect which was blocked when a calm mother was present during shock exposure. No sex differences in the effect of maternal presence on the stress response to shock and no interactions between sex and prenatal drug exposure were observed. Although VPA-exposed pups may show impaired responsivity to maternal cues in early infancy, maternal presence is still capable of regulating the stress response in VPA-exposed pups. In this study we demonstrate the importance of utilizing multiple batteries of tests in assessing behavior, dissecting the behavior on one test into different components. Our results inform about the underlying behavioral characteristics of some of the ASD phenotypes, including sex differences reported by clinical studies, and could shed light on potential opportunities for intervention.

## 1. Introduction

The prevalence of children diagnosed with autism spectrum disorders (ASDs) has steadily increased over the past 15 years (Baio et al., 2018). Disrupted processing of social cues and altered social behaviors are among the core symptoms of ASDs (Lord et al., 2018). Most children with ASDs are diagnosed after age 3 (Sheldrick et al., 2017); however, children who will later be diagnosed with an ASD show differences in visual processing and attentiveness to social cues as early as the first year of life (Zwaigenbaum et al., 2005). Specifically, poor visual attention, stronger negative emotional displays, and delayed verbal abilities were observed in infants that were later diagnosed with autism spectrum disorder (Zwaigenbaum et al., 2005).

These sensory abilities that may be altered or develop later in ASDs are critical for the process of attachment learning. Attachment learning refers to the process by which infants learn which sensory cues are associated with the caregiver and enables the infant to seek out parental care when under threat or stressed (Bowlby, 1977). The presence of a caregiver can suppress a child’s stress response (Hostinar et al., 2015; Nachmias et al., 1996) and the quality of that attachment bond modulates the effectiveness of this stress buffering effect (Gunnar, 2017). It is not yet known whether caregiver buffering of stress is intact in children with an ASD or infants that will later be diagnosed with an ASD.

Humans exposed to the epilepsy drug valproic acid (VPA) during pregnancy have higher rates of autism (Christensen et al., 2013); therefore, exposing prenatal laboratory rodents to VPA and examining the resulting phenotypes has been one approach to modeling aspects of ASDs in rodents (Dufour-Rainfray et al., 2011). As adolescents, rodents exposed to VPA during pregnancy show reduced social play behavior (Markram et al., 2008; Schneider & Przewłocki, 2005), social exploration (Markram et al., 2008; Schneider & Przewłocki, 2005), and social interactions as juveniles and adults (Kim et al., 2014). For example, adolescent male and female rats exposed to VPA in utero approached a conspecific less frequently than those exposed to saline in utero (Barrett et al., 2017).

Relatively less is known about how VPA-treated rats respond to social cues in infancy. Laboratory rat infants, like human infants, undergo the process of attachment learning. Olfactory cues associated with the mother and nest become powerful motivators of infant behavior and maintain the attachment bond (Debiec & Sullivan, 2017). Upon separation from the mother, a rat pup will approach olfactory stimuli associated with the mother and/or nest (Bolles & Woods, 1964; Mendez-Gallardo & Robinson, 2014) and will vocalize to obtain the mother’s attention and care (Brunelli et al., 1994; Hofer & Shair, 1978). Infant rats exposed to prenatal VPA appear to have deficits in this attachment behavior. At postnatal day 9, infant rats exposed to prenatal VPA were slower to approach home cage bedding than infant rats exposed to prenatal saline (Schneider & Przewłocki, 2005). This delayed approach behavior was absent in VPA-treated pups at postnatal day 11 and 13 (Schneider & Przewłocki, 2005). A more complex analysis of how infant rats exposed to prenatal VPA respond to maternal cues has yet to be performed.

Furthermore, it is unclear if other vital attachment-associated behaviors, such as maternal buffering of stress, are impaired in VPA-exposed pups. As in human infants, infant rats rely on maternal regulation of the stress response axis. Maternal presence, direct maternal care, and cues associated with the mother blunt or entirely suppress the infant rat’s stress response to various stressors including predator odor and shock (Moriceau & Sullivan, 2006; Shionoya et al., 2007; Stanton et al., 1987; Stanton & Levine, 1990; Suchecki et al., 1993; Wiedenmayer et al., 2003).

Understanding when and how ASD features emerge could enable early identification of at-risk individuals and early intervention. Indeed, training parents of infants at high risk of autism to pay closer attention to their infants’ communication style and adjust accordingly was associated with better outcomes months later (Green et al., 2017). Here, we first examined the behavioral response to social odors in infant rats exposed to VPA or a control saline solution at E12.5. Next, we asked whether maternal regulation of the stress response was intact in infant rats exposed to VPA relative to those exposed to saline during pregnancy. Because there are sex differences in the diagnosis and symptomology of ASDs (Ferri et al., 2018), we also looked for potential sex differences in these pups with ASD-like features

## 2. Methods

### 2.1 Animals

Outbred female and male Sprague-Dawley rats of breeding age were obtained from Charles River. The room was set to a 12:12 h light:dark cycle and maintained at 22 ± 2 °C. Fod and water were freely available. Estrus cycle stage was determined by a vaginal lavage performed in the late afternoon and then examining cytology under a light microscope. Sexually receptive females were paired with a male rat overnight and separated the following morning.

Valproic acid (VPA) (Sigma) was dissolved in sterile saline to obtain a 250 mg/mL solution. At embryonic day 12.5, pregnant females received an IP injection of a 500 mg/kg dose of VPA or an equivalent volume of sterile saline (Schneider & Przewłocki, 2005). The day after birth (postnatal day 1 (P1)) litters were culled to a maximum of 12 pups/litter. Mothers and pups were subsequently left undisturbed until the day of testing. Experiments were performed in the light phase by a female experimenter. All procedures were approved by the University of Michigan Institutional Animal Care & Use Committee.

### 2.2 Odor Preference Test (OPT)

In early infancy (postnatal day (P) 6-7) VPA-exposed and saline-exposed pups underwent an odor preference test (OPT) for social odors. VPA-exposed pups (*n* = 12 females, 11 males) and saline-exposed pups (*n* = 11 females, 13 males) were retrieved from the home cage 3 at a time and brought to a separate room from the colony room. The OPT took place in a 20.8 cm x 31 cm x 17.7 cm (width x length x height) arena with a wire mesh floor over a base that contained clean bedding at one end (20.8 cm x 7.5 cm) and contained soiled bedding laden with social olfactory cues from the pup’s homecage at the other end (20.8 cm x 7.5 cm). The test consisted of 5 1-minute trials. Pups were run one at a time; the pups that were not currently completing the OPT were kept on the other side of the room in a cage on top of a heating pad. At the beginning of each trial, the pup was placed in the center of the arena. In order to alleviate the development of a side preference, the orientation of the pup was alternated each trial and the entire arena was rotated after 2 or 3 trials. In mid-infancy (P13), separate groups of VPA and saline-exposed pups underwent an OPT. VPA-exposed pups (n = 9 females, 7 males) and saline-exposed pups (n = 9 females, 9 males) underwent a test for social odor preference. Video analysis was performed in Ethovision. Each pups’ latency to enter each zone of the arena, the number of entries into each zone of the arena, the amount of time spent in each zone of the arena, and total distance traveled during the test was recorded. A pup was defined as being in a zone if the center-point of the pup was inside the zone.

### 2.3 Maternal buffering of stress response experiments

On PND 6 or 7, VPA and saline-treated pups were placed in an empty cage lined with an absorbent pad. No more than 6 pups underwent the stress procedure at once. A heating pad was placed underneath the cage in order to maintain pups’ body temperature. After 10 minutes of habituation, pups were exposed to a 0.5 mA shock to the tail. Pups received 10 additional tail shocks each separated by a 4-minute inter-trial interval. Immediately following stress exposure, pups were brought one at a time into an adjacent room and sacrificed. Baseline corticosterone levels for VPA-treated animals and saline-treated animals were measured by collecting trunk blood from separate groups of animals that did not undergo stress exposure and were instead euthanized immediately after removal from the homecage. Trunk blood was collected in EDTA tubes and centrifuged at 3000 rpm for 10 minutes at 4 C. Following centrifugation, serum was extracted and stored at −80 C. All samples were analyzed in a single radioimmunoassay performed at the UM Core Facility.

### 2.4 Statistical Analysis

Data were analyzed utilizing two-way ANOVAs and unpaired t-tests using GraphPad Prism. Tukey’s post-hoc test was used where appropriate.

## 3. Results

Given that the initial group to develop the VPA-exposure model in rat pups (Schneider & Przewłocki, 2005) found that VPA pups had lower mean weights than saline-treated pups, we measured and compared weights in a subset of pups from multiple litters. At P6-7, the average weight of VPA-exposed pups (*n* = 48) was 15.59 grams whereas the average weight of saline-exposed pups (*n* = 87) was 16.41 grams. These differences were not significant, *t*(133) = 1.89, *p* = 0.06. In P13 pups, VPA-exposed pups (*n* = 10) had an average weight of 23.60 grams and saline-exposed pups (*n* = 18) had an average weight of 28.74 grams. These differences were significant *t*(26) = 3.75, *p* = 0.0009.

### 3.1 P6-7 Odor Preference Test

VPA-treated and saline-treated animals traveled similar distances during the OPT, *t*(33) = 1.34, *p* = 0.19 (Figure 1D). However, striking differences began to emerge once we looked at how VPA-treated and saline-treated animals responded to social cues during the test. A two-way ANOVA revealed a significant main effect of bedding type (*F*(1,90) = 55.17, *p* < 0.0001) and drug treatment (*F*(1,90) = 12.21, *p* = 0.0007) on latency to enter a zone of the odor preference arena, but no significant interaction (*F*(1,90) = 1.28, *p* = 0.26) (Figure 1A). Both VPA and saline-treated pups showed a shorter latency to enter the home cage bedding zone of the odor preference arena (*p* < 0.0001, *p* = 0.0001). VPA-treated pups showed a longer latency to enter the clean bedding zone of the odor preference arena than saline-treated pups (*p* = 0.0081) but not the home cage bedding zone (*p* = 0.35). This may suggest that VPA-treated pups have reduced motivation to explore the arena than saline-treated pups.

**Figure 1.**
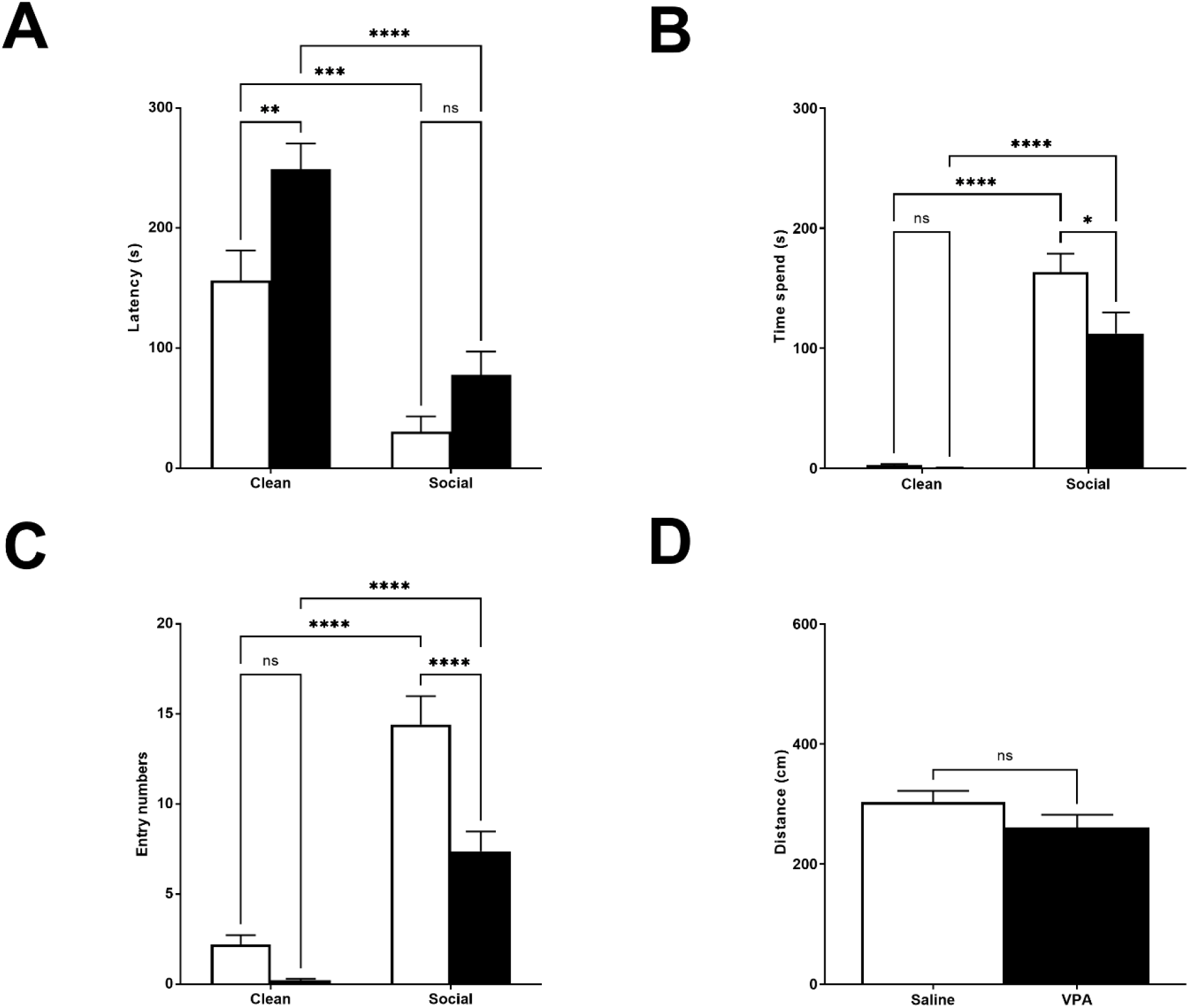
*A.) P6-7: Latency to Enter Zones*. We observed significant main effects of bedding type, *p* < 0.0001, and drug exposure, *p* = 0.007. VPA-exposed pups showed a longer latency to enter the clean bedding zone, *p* = 0.008. *B.) P6-7: Time Spent in Zones*. We observed significant main effects of bedding type, *p* < 0.0001, drug exposure, *p* < 0.0001, and a significant interaction, *p* = 0.014. VPA-exposed pups spent significantly less time in the SB zone, *p* < 0.0001. *C.) P6-7: Number of Zone Entries*. We observed significant main effects of bedding type, *p* < 0.04, drug exposure, *p* < 0.0001, and a significant interaction, *p* = 0.02. VPA-exposed pups made significantly fewer entries into the social bedding zone, *p* = 0.01. *D.) P6-7: Distance Traveled during Test*. VPA-exposed pups did not differ from saline-exposed pups in their distance traveled during the test.

When examining the amount of time spent in each zone of the odor preference arena, a two-way ANOVA revealed a significant main effect of bedding type (*F*(1,90) = 138.4, *p* < 0.0001), drug treatment (*F*(1,90) = 5.32, *p* = 0.02), and a significant interaction (*F*(1,90) = 4.42, *p* = 0.04) (Figure 1B). Although both groups showed a preference for the home cage bedding relative to the clean bedding (*p*’s < 0.0001), VPA-exposed pups spent significantly less time in the home cage bedding zone, (*p* = 0.01).

A two-way ANOVA revealed a significant main effect of bedding type (*F*(1,90) = 93.34, *p* < 0.0001) and drug treatment (*F*(1,90) = 20.20, *p* < 0.0001) on the number of entries into a zone of the odor preference arena, and a significant interaction (*F*(1,90) = 6.30, *p* = 0.01) (Figure 1C). Both VPA and saline-treated pups made more entries into the home cage bedding zone of the odor preference arena (*p*’s < 0.0001). However, VPA-treated pups made fewer entries into the home cage bedding zone of the odor preference arena than saline-treated pups (*p* < 0.0001). This could suggest that VPA-treated pups have reduced motivation to approach the home cage bedding than saline-treated pups, or that VPA-treated pups are less motivated to explore in general than saline-treated pups.

### 3.2 P6-7 Odor Preference Test: Sex Differences

Next, we analyzed differences between VPA and saline-treated animals separately within female and male animals. We observed no effect of sex on total distance traveled during the OPT (*F*(1,54) = 0.004, *p* = 0.95), no effect of drug exposure (*F*(1,54) = 1.22, *p* = 0.27), and no interaction between sex and drug exposure (*F*(1,54) = 0.04, *p* = 0.84).

In female pups, a two-way ANOVA revealed a significant main effect of bedding type (*F*(1,42) = 26.35, *p* < 0.0001) on latency to enter a zone of the odor preference arena, but no effect of drug treatment (*F*(1,42) = 1.23, *p* = 0.27) or significant interaction (*F*(1,42) = 0.03, *p* = 0.86) (Figure 2A). Both female VPA and female saline-treated pups showed a shorter latency to enter the home cage bedding zone of the odor preference arena (*p* = 0.0023, *p* = 0.0071). Female VPA-treated pups did not show a longer latency to enter the familiar odor zone of the odor preference arena than saline-treated pups (*p* = 0.91) or the home cage bedding zone (*p* = 0.80). In male animals, we observed a different pattern of results. A two-way ANOVA revealed a significant main effect of bedding type (*F*(1,44) = 29.88, *p* < 0.0001), a significant main effect of drug treatment (*F*(1,44) = 15.07, *p* = 0.0003), but no significant interaction (*F*(1,44) = 2.14, *p* = 0.15) (Figure 2B). Both male VPA-treated pups (*p* = 0.0001) and male saline-treated pups (*p* = 0.02) showed a shorter latency to enter the social odor zone of the odor preference arena. Male VPA-treated pups and male saline-treated pups did not differ in their latency to enter the familiar odor zone of the odor preference arena (*p* = 0.33) however male VPA-treated pups showed a higher latency to enter the clean bedding zone of the odor preference arena than male saline-treated pups (*p* = 0.003).

**Figure 2.**
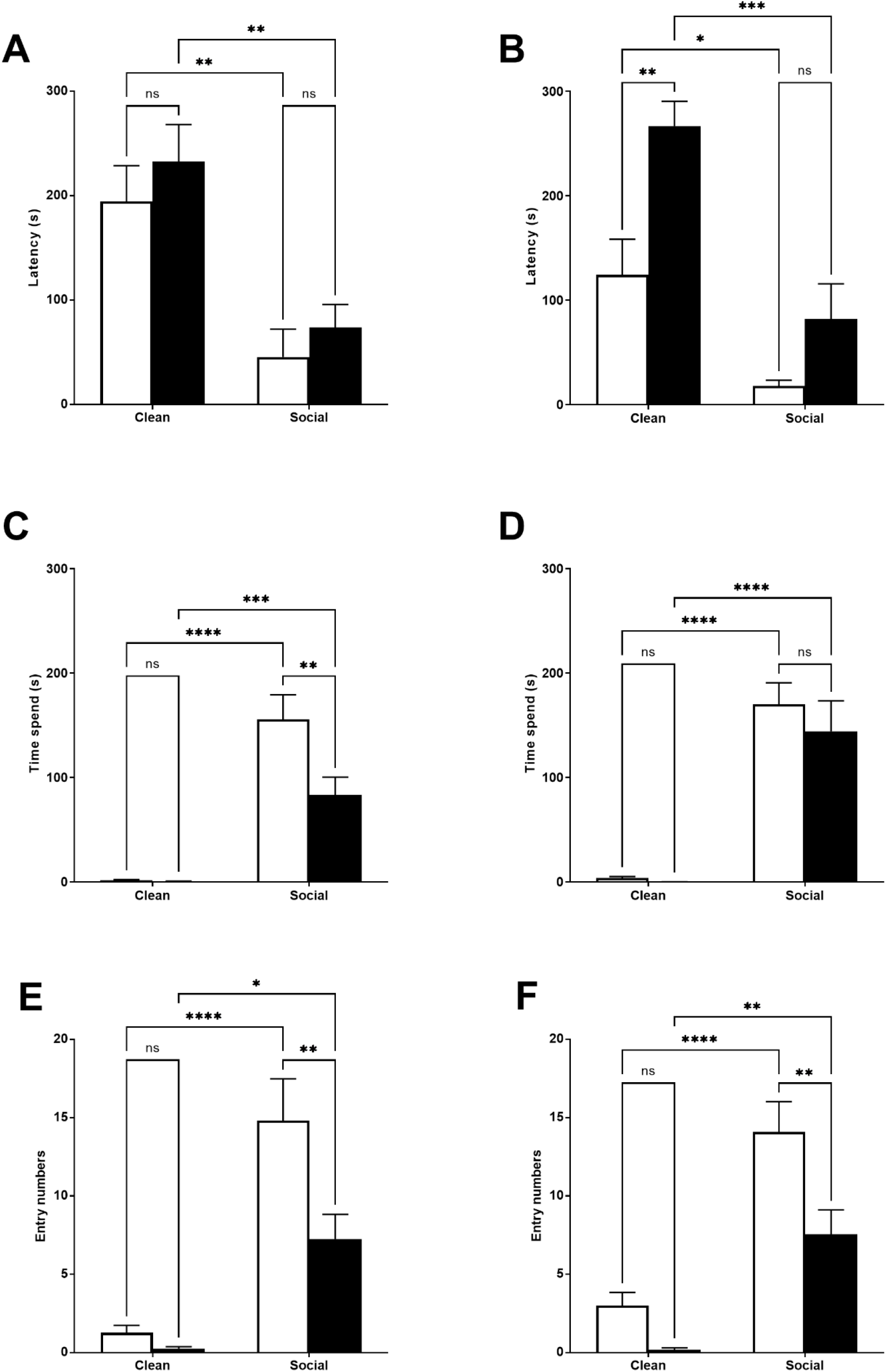
*A-B.) P6-7 Sex Differences: Latency to Enter Zones*. We observed a significant main effect of bedding type in females (A), *p* < 0.0001, and males (B), *p* < 0.0001. There was a significant main effect of drug exposure in males alone, *p* = 0.003. *C-D.) P6-7 Sex Differences: Time Spent in Zones*. We observed a significant main effect of bedding in both females (C), p < 0.0001, and males (D), p < 0.0001. In females, we also observed a significant main effect of drug exposure, *p* = 0.015, and a significant interaction, *p* = 0.02. Female VPA-exposed pups spent significantly less time in the SB zone than female saline-exposure pups, *p* = 0.005. *E-F.) P6-7 Sex Differences: Number of Zone Entries*. We observed significant main effects of bedding type and drug exposure in females (E), *p* < 0.0001 and *p* = 0.008, and males (F), *p* < 0.0001 and *p* = 0.0013. There was also a significant interaction in females, *p* = 0.04. Both female, *p* = 0.006, and male, *p* = 0.0075, VPA-exposed pups made significantly fewer entries into the SB zone.

We next examined whether our drug treatment group differences observed on time spent in different zones of the odor preference arena differed by sex. In female animals, a two-way ANOVA revealed a significant main effect of bedding type (*F*(1,42) = 66.79, *p* < 0.0001) and drug treatment (*F*(1,42) = 6.46, *p* = 0.015) and a significant interaction effect (*F*(1,42) = 6.09, *p* = 0.02) on the amount of time spent in each zone of the odor preference arena (Figure 2C). Both VPA-treated female animals (*p* = 0.001) and saline-treated female animals (*p* < 0.0001) spent more time in the familiar odor zone of the odor preference arena than in the clean bedding zone. However, VPA-treated female animals spent significantly less time in the familiar odor zone of the odor preference arena than saline-treated female animals (*p* = 0.005). In male animals, we observed a significant main effect of bedding type on the amount of time spent in each zone of the odor preference arena (*F*(1,44) = 78.8, *p* < 0.0001) but no significant main effect of drug treatment (*F*(1,44) = 0.70, p = 0.41) or significant interaction effect (*F*(1,44) = 0.41, p = 0.53) (Figure 2D). Both male VPA-treated animals (*p* < 0.0001) and male saline-treated animals (*p* < 0.0001) spent more time in the social odor zone of the odor preference arena than in the clean zone of the odor preference arena. However, male VPA-treated and male saline-treated animals did not differ in the amount of time they spent in the social odor zone of the odor preference arena (*p* = 0.72).

Finally, we examined whether we observed different group effects in zone entry numbers separately in female and male animals. In female animals, a two-way ANOVA revealed a significant main effect of bedding type (*F*(1,42) = 44.61, *p* < 0.0001) and drug treatment (*F*(1,42) = 7.80, p = 0.008) on the number of entries into a zone of the odor preference arena, and a significant interaction (*F*(1,42) = 4.53, *p* = 0.04) (Figure 2E). Both female VPA and female saline-treated pups made more entries into the home cage bedding zone of the odor preference arena (*p* = 0.01, *p* < 0.0001). VPA-treated female pups made fewer entries into the home cage bedding zone of the odor preference arena than saline-treated female pups (*p* = 0.006) but not the clean bedding zone (*p* = 0.97). In male animals, a two-way ANOVA revealed significant main effects of bedding (*F*(1,44) = 46.18, *p* < 0.0001) and drug treatment (*F*(1,44) = 11.87, *p* = 0.001), but no significant interaction effect (*F*(1,44) = 1.87, *p* = 0.18) (Figure 2F). Both VPA-treated (*p* = 0.003) and saline-treated (*p* < 0.0001) male animals made more entries into the social odor zone of the odor preference arena. However, male saline-treated animals made more entries into the familiar odor zone of the odor preference arena than male VPA-treated animals (*p* = 0.008).

### 3.3 P13 Odor Preference Test

We next examined VPA-treated and saline-treated pups’ behavior in the OPT in a cohort of P13 animals. As observed in younger pups, P13 VPA and saline-treated pups traveled similar differences during the OPT (*t*(20) = 0.48, *p* = 0.64) (Figure 3D). There was a significant main effect of bedding type on latency to enter a zone (*F*(1,64) = 80.04, *p* < 0.0001) but no significant effect of drug treatment (*F*(1,64) = 0.19, *p* = 0.66) and no significant interaction (*F*(1,64) = 0.43, *p* = 0.52) (Figure 3A). Both VPA and saline-treated pups showed a shorter latency to enter the social bedding zone relative to the clean bedding zone (*p*’s < 0.0001) and did not differ in their latency to enter the social bedding zone (*p* = 1.00).

**Figure 3.**
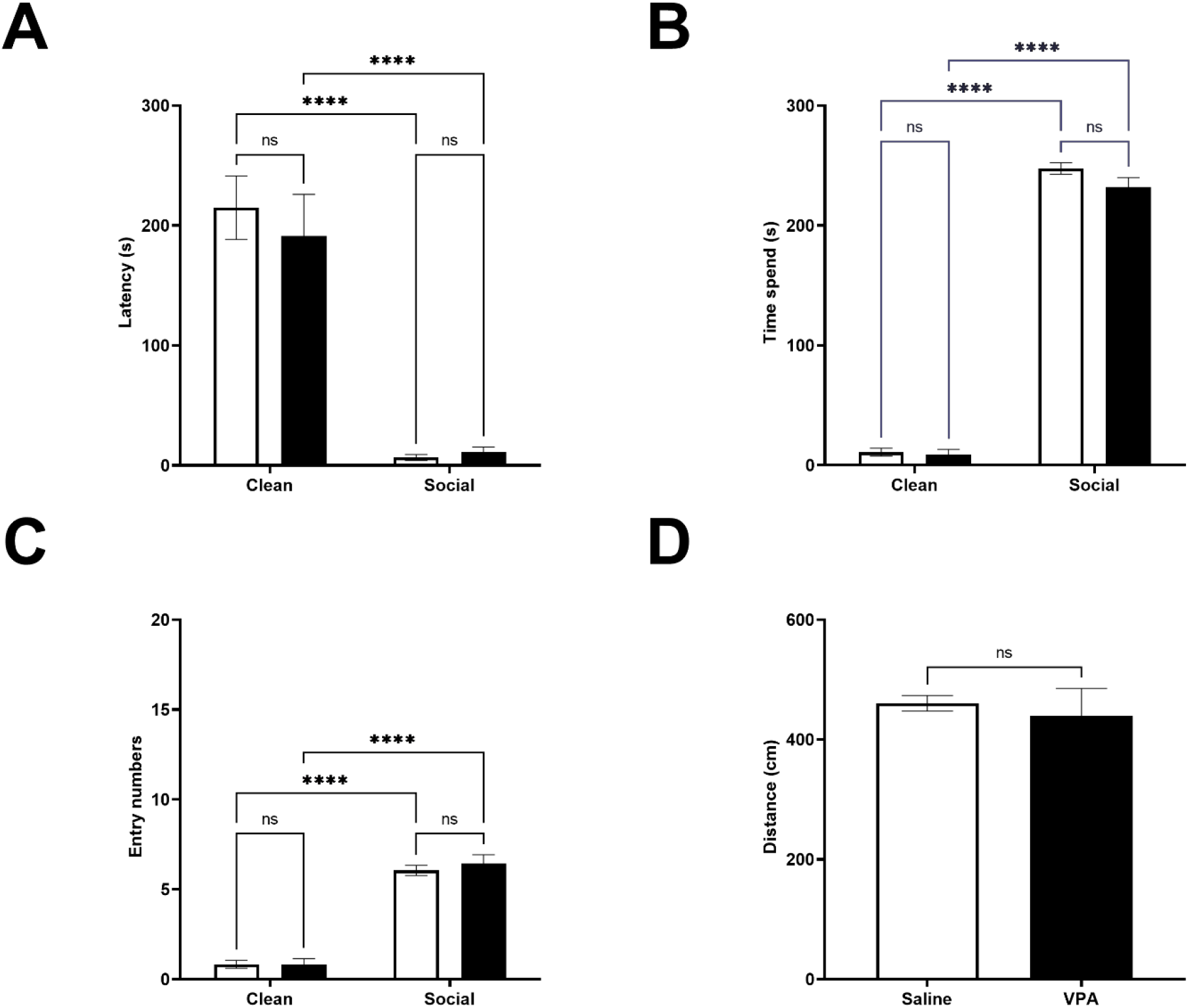
*A.) P13: Latency to Enter Zones*. We observed a significant main effect of bedding type, *p* < 0.0001. Both VPA-exposed and saline exposed-pups showed a shorter latency to enter the social bedding zone than the clean bedding zone, *p’s* < 0.0001. *B.) P13: Time Spent in Zones*. We observed a significant main effects of bedding type, *p* < 0.0001. Both VPA-exposed pups and saline-exposed pups spent significantly more time in the social bedding zone than the clean bedding zone, *p’s* < 0.0001. *C.) P13: Number of Zone Entries*. We observed a significant main effects of bedding type, *p* < 0.00001. VPA-exposed pups and saline-exposed pups made significantly fewer entries into the social bedding zone relative to the clean bedding zone *p’s* < 0.00001. *D.) P13: Distance Traveled during Test*. VPA-exposed pups did not differ from saline-exposed pups in their distance traveled during the test, *p* = 0.64.

When examining the time spent in each zone of the OPT, we found a significant main effect of bedding type (*F*(1,64) = 1880, *p* < 0.0001), but no significant effects of drug treatment (*F*(1,64) = 2.71, *p* = 0.10) or interaction (*F*(1,64) = 1.66, *p* = 0.20) (Figure 3B). Tukey’s multiple comparisons test revealed that VPA and saline-treated pups spent more time in the social bedding zone than the clean bedding zone (*p*’s < 0.0001). VPA and saline-treated pups spent similar amounts of time in the social bedding zone during the test (*p* = 0.17).

Similar effects were observed when investigating how many entries pups made into the zones of the OPT. There was a significant main effect of bedding type (*F*(1,64) = 261.8, *p* < 0.0001), but again no significant main effect of drug treatment (*F*(1,64) = 0.29, *p* = 0.59) and no significant interaction (*F*(1,64) = 0.36, *p* = 0.55) (Figure 3C). VPA and saline-treated pups made more entries into the social bedding zone relative to the clean bedding zone (*p*’s < 0.0001), but there were no differences between VPA and saline-treated pups in the number of entries made into the social bedding zone (*p* = 0.85).

### 3.4 P13 Odor Preference Test: Sex Differences

Given that we observed some sex differences in behavior during the OPT at P6-7, we next asked whether we observed similar differences at P13. We did not find significant effects of sex (*F*(1,18) = 0.46, *p* = 0.51), drug exposure (*F*(1,18) = 0.21, *p* = 0.65) or interaction between sex and drug exposure (*F*(1,18) = 0.003, *p* = 0.96) on distance traveled during the OPT.

In female pups, there was a significant main effect of bedding type (*F*(1,32) = 55.63, *p* < 0.0001) but no significant main effect of drug exposure (*F*(1,32) = 0.08, *p* = 0.77) or significant interaction effect (*F*(1,32) = 0.00007, *p* = 0.99) on latency to enter the zones of the OPT (Figure 4A). All female pups were quicker to enter the home cage bedding zone than the clean bedding zone (*p*’s < 0.0001) and VPA-treated and saline-treated female pups did not differ in their latency to enter the home cage bedding zone (*p* = 1.00). This pattern of results was also observed in male pups. We observed a significant main effect of bedding type on latency to enter the zones of the OPT (*F*(1,28) = 25.79, p < 0.0001), but no significant main effect of drug exposure (*F*(1,28) = 1.02, *p* = 0.32) and no significant interaction (*F*(1,28) = 1.07, *p* = 0.31) (Figure 4B). Both VPA-treated and saline-treated male pups showed a shorter latency to enter the social bedding zone than the clean bedding zone (*p* = 0.05, *p* = 0.0004), and VPA-treated and saline-treated male pups showed similar latencies to enter the social bedding zone (*p* = 0.99).

**Figure 4.**
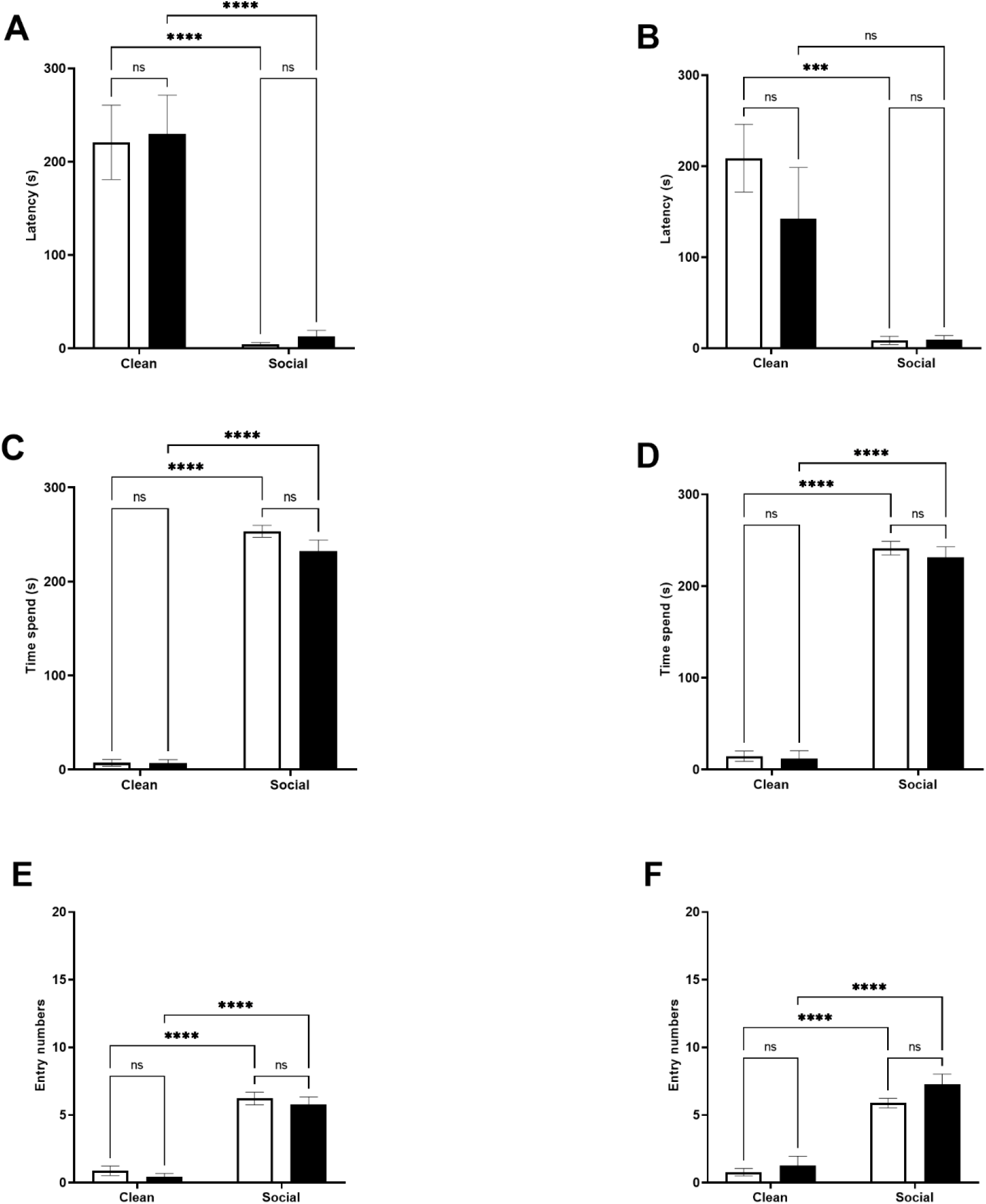
*A-B.) P13 Sex Differences: Latency to Enter Zones*. We observed a significant main effect of bedding type in females (A), *p* < 0.0001, and males (B), *p* < 0.0001. *C-D.) P13 Sex Differences: Time Spent in Zones*. We observed a significant main effect of bedding in both females (C), p < 0.0001, and males (D), p < 0.0001. *E-F.) P13 Sex Differences: Number of Zone Entries*. We observed a significant main effect of bedding in both females (C), p < 0.0001, and males (D), p < 0.0001.

When examining the amount of time female pups spent in each zone of the OPT, we found a significant main effect of bedding type (*F*(1,32) = 1111, *p* < 0.0001), but no main effect of drug exposure (*F*(1,32) = 2.32, *p* = 0.14) or interaction effect (*F*(1,32) = 2.03, *p* = 0.16 (Figure 4C). Female pups spent more time in the social bedding zone regardless of prenatal treatment (*p*’s < 0.0001) and both female VPA-treated and saline-treated pups spent a similar amount of time in the social bedding zone (*p* = 0.18). In male pups, we found a significant main effect of bedding type (*F*(1,28) = 731.3, *p* < 0.0001), but no significant main effect of drug exposure (*F*(1,28) = 0.60, *p* = 0.44) or interaction effect (*F*(1,28) = 0.24, *p* = 0.63) on time spent in the zones of the OPT (Figure 4D). Male VPA-exposed and saline-exposed pups spent more time in the social bedding zone than the clean bedding zone (*p*’s < 0.0001) and did not differ from each other in time spent in the social bedding zone (*p* = 0.81).

Finally, in female pups we observed a significant main effect of bedding type (*F*(1,32) = 156.9, *p* < 0.0001), but no main effect of drug exposure (*F*(1,32) = 1.09, *p* = 0.30) or interaction (*F*(1,32) = 0.00, *p* > 0.99) on entries made into the zones of the OPT (Figure 4E). Both VPA and saline-treated pups made more entries into the social bedding zone (*p*’s < 0.0001) and did not differ from each other in the number of entries made into the social bedding zone (*p* = 0.88). In male pups, we observed a significant main effect of bedding type (*F*(1,28) = 118.8, *p* < 0.0001), but no main effect of drug exposure (*F*(1,28) = 3.49, *p* = 0.07) or interaction effect (*F*(1,28) = 0.76, *p* = 0.39) (Figure 4F). Both VPA-treated and saline-treated male pups entered the social bedding zone more frequently than the clean bedding zone (*p*’s < 0.0001) and they did not differ from each other in the number of entries made into the social bedding zone (*p* = 0.24).

### 3.5 Maternal Buffering of Stress Response

We next investigated whether maternal presence was equally effective in regulating the corticosterone response to shock in saline and VPA-treated pups at P6-7. VPA-exposed pups and saline-exposed pups did not differ in their weight on test day (*t*(98) = 0.98, *p* = 0.33). A 2 x 3 ANOVA revealed a significant main effect of experimental condition, (*F*(2,93) = 81.77, *p* < 0.0001) (Figure 5A). There was no significant main effect of prenatal treatment (*F*(1,93) = 0.46, *p* = 0.50), and no significant interaction effect (*F*(2,93) = 0.86, *p* = 0.43). Tukey’s multiple comparisons test showed that both saline and VPA-treated animals exposed to shock showed significantly higher corticosterone levels than baseline groups (*p*’s < 0.0001) and the groups exposed to shock in the presence of a calm mother (*p*’s < 0.0001). Corticosterone levels in saline (*p* = 0.28) and VPA-treated pups (*p* = 0.95) shocked in the presence of a calm mother were not significantly different from baseline corticosterone levels.

**Figure 5.**
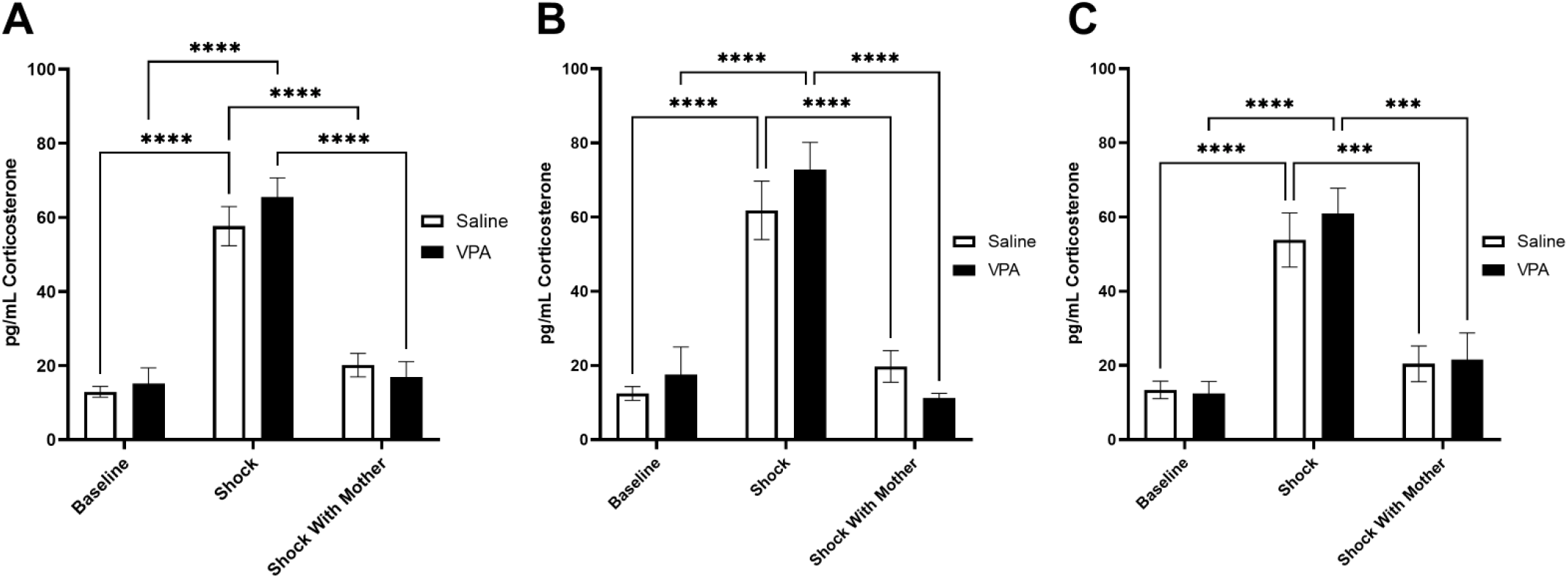
*A.) Maternal Buffering of Stress Response to Shock at P6-7*. We observed a significant main effect of experimental condition, *p* < 0.0001 on pup corticosterone levels. Both saline and VPA-treated pups exposed to shock showed significantly higher corticosterone levels than baseline groups (*p*’s < 0.0001) and the groups exposed to shock in the presence of a calm mother (*p*’s < 0.0001). *B.) Maternal Buffering of Stress Response to Shock in Females at P6-7*. We observed a significant main effect of experimental condition, *p* < 0.0001 on female pup corticosterone levels. Both saline and VPA-treated female pups exposed to shock showed significantly higher corticosterone levels than baseline groups (*p*’s < 0.0001) and the groups exposed to shock in the presence of a calm mother (*p*’s < 0.0001). *C.) Maternal Buffering of Stress Response to Shock in Males at P6-7*. We observed a significant main effect of experimental condition, *p* < 0.0001 on male pup corticosterone levels. Both saline and VPA-treated male pups exposed to shock showed significantly higher corticosterone levels than baseline groups (*p*’s < 0.0001) and the groups exposed to shock in the presence of a calm mother (saline *p* = 0.0001, VPA *p* = 0.0003).

Similar results were observed when we split our data by sex. Within female pups, there was a significant main effect of experimental condition on corticosterone levels (*F*(2,42) = 49.11, *p* < 0.0001) but no significant main effect of prenatal treatment (*F*(1,42) = 0.27, *p* = 0.61), or interaction effect (*F*(2,42) = 1.28, *p* = 0.29) (Figure 5B). Within both saline and VPA-treated female pups, corticosterone levels were significantly higher in shocked pups relative to baseline corticosterone levels and relative to levels in pups shocked with a calm mother present (*p*’s < 0.0001). Corticosterone levels did not differ between female pups taken directly from the home cage and female pups shocked with a calm mother present (saline *p* = 0.56, VPA *p* = 0.78).

In male pups, a 2 x 3 ANOVA found a significant main effect of experimental group (*F*(2,45) = 33.44, *p* < 0.0001) but no significant main effect of prenatal treatment (*F*(1,45) = 0.25, *p* = 0.62), and no significant interaction effect (*F*(2,45) = 0.27, p = 0.77) (Figure 5C). In saline and VPA-treated male pups, corticosterone levels in the shock group were significantly higher than baseline corticosterone levels (*p*’s < 0.0001) and significantly higher than in male pups shocked with a calm mother present (saline *p* = 0.0001, VPA *p* = 0.0003). Corticosterone levels did not differ between those in the baseline groups and those in the shock with calm mother groups (saline *p* = 0.63, VPA *p* = 0.61).

## 4. Discussion

Here, we found that pups treated with VPA during pregnancy may have impaired social recognition and/or may be less motivated to approach social odors in early infancy. In the OPT, 6-7 day old VPA-exposed pups spent significantly less time in proximity with the home cage bedding and made fewer entries into the zone containing home cage bedding. These effects were particularly prominent in female pups. At postnatal day 13, VPA-exposed pups and saline-exposed pups showed similar preferences for home cage bedding. Although VPA-exposed pups may initially have a deficit in this attachment-related behavior they do recover typical responses to home cage bedding in later infancy. There may be a developmental lag in the pup’s ability to learn to associate olfactory cues with the mother or, alternatively, there may be a developmental lag in the pup’s ability to respond appropriately to olfactory cues associated with the mother.

Our findings parallel, in part, what has been previously shown in the VPA autism-like phenotype. In a similar OPT, Schneider & Przewłocki (Schneider & Przewłocki, 2005) found that P9 VPA-exposed rats had a higher latency to enter the zone containing home cage bedding than saline-exposed rats, an effect which was no longer present in P11 and P13 rats. Time spent in each zone of the OPT and frequency of entries into each zone of the OPT were not recorded, and it is in these behaviors that we observed differences between P6-7 VPA-treated and saline-treated pups in our study. Development progresses rapidly in infant rats; although P6-7 and P9 pups are still at an age in which they are spending the majority of their time in the nest (Bolles & Woods, 1964), P9 pups are more accomplished at crawling and move faster than P6-7 pups (Altman & Sudarshan, 1975). Entry latency could be a measure more sensitive to motor development than number of entries and time spent and thus differences between VPA-exposed and saline-exposed pups may be more difficult to observe.

To the best of our knowledge, this study is the first to look for potential sex differences in pup behavior in the VPA model. In both P6-7 male and female pups, we found that VPA-exposed pups made fewer entries into the social bedding zone of the OPT and no difference between VPA-exposed and saline-exposed pups in the latency to enter the social bedding zone of the OPT. Interestingly, female VPA-exposed pups spent less time in the social bedding zone of the OPT than saline-exposed pups but male VPA-exposed pups did not differ from male saline-exposed pups in this behavior. Rat mothers tend to provide more care to their male pups than their female pups (Deviterne & Desor, 1990; Moore & Morelli, 1979; Richmond & Sachs, 1984), so perhaps this provides male pups with extra reinforcement of attachment learning and enables VPA-exposed male pups to recover a more typical response to maternal cues. Some findings from adult rats exposed to VPA in utero suggest that the effects may manifest differently in male and female animals (Barrett et al., 2017), so this may also explain the difference we observed here. In P13 pups, we noted no differences between VPA and saline-exposed pups in the OPT and these results did not change when we divided animals by sex.

To address whether another vital function which relies on detection of social cues was altered in VPA-exposed pups, we measured infant rats’ stress response to repeated shocks with and without maternal presence. Both saline and VPA-exposed pups showed robust corticosterone responses to repeated shocks, an effect which was blocked when a calm mother was present during shock exposure. No sex differences in the effect of maternal presence on the stress response to shock and no interactions between sex and prenatal drug exposure were observed. These data indicate that although VPA-exposed pups may show impaired responsivity to maternal cues in early infancy, maternal presence is still capable of regulating the stress response in VPA-exposed pups. This early alteration of social odor-motivated behaviors may be particularly important because infant rat attachment learning relies on olfactory social cues. Furthermore, because VPA-exposed pups show deficits in approaching maternal cues, they may experience more stressful events in the absence of the mother. This increase in early life stress could explain, in part, the increased anxiety-like behavior observed in adult animals treated with VPA during prenatal development (Barrett et al., 2017; Ellenbroek et al., 2016; Li et al., 2017; Markram et al., 2008).

Children who develop ASDs often begin to show social differences beginning in infancy (Zwaigenbaum et al., 2005) and may benefit from early intervention (Green et al., 2017). Identifying these social differences and investigating how they may affect vital attachment functions such as maternal buffering of stress is critical for improving outcomes for these individuals. Animal models, such as the VPA model for ASDs, can provide the critical initial steps in addressing some of these questions. In this study we demonstrate the importance of utilizing multiple batteries of tests in assessing behavior, dissecting the behavior on one test into different components. Our results inform about the underlying behavioral characteristics of some of the ASD phenotypes, including sex differences reported by clinical studies, and could shed light on potential opportunities for intervention.

## Acknowledgements

This research was supported by the K08 MH014743-01A1 and NARSAD Young Investigator Grant from the Brain & Behavior Research Foundation to JD.

## References

Altman, J., & Sudarshan, K. (1975). Postnatal development of locomotion in the laboratory rat. Animal Behaviour, 23, 896–920. https://doi.org/ https://doi.org/10.1016/0003-3472(75)90114-1

Baio, J., Wiggins, L., Christensen, D. L., Maenner, M. J., Daniels, J., Warren, Z., Kurzius-Spencer, M., Zahorodny, W., Rosenberg, C. R., White, T., Durkin, M. S., Imm, P., Nikolaou, L., Yeargin-Allsopp, M., Lee, L. C., Harrington, R., Lopez, M., Fitzgerald, R. T., Hewitt, A., … Dowling, N. F. (2018). Prevalence of autism spectrum disorder among children aged 8 Years - Autism and developmental disabilities monitoring network, 11 Sites, United States, 2014. MMWR Surveillance Summaries, 67(6). https://doi.org/10.15585/mmwr.ss6706a1

Barrett, C. E., Hennessey, T. M., Gordon, K. M., Ryan, S. J., McNair, M. L., Ressler, K. J., & Rainnie, D. G. (2017). Developmental disruption of amygdala transcriptome and socioemotional behavior in rats exposed to valproic acid prenatally. Molecular Autism, 8(1), 1–17. https://doi.org/10.1186/s13229-017-0160-x

Bolles, R. C., & Woods, P. J. (1964). The ontogeny of behaviour in the albino rat. Animal Behaviour, 12(4), 427–441. https://doi.org/10.1016/0003-3472(64)90062-4

Bowlby, J. (1977). The Making and Breaking of Affectional Bonds: I. Aetiology and Psychopathology in the Light of Attachment Theory. British Journal of Psychiatry, 130(3), 201–210. https://doi.org/ DOI: 10.1192/bjp.130.3.201

Brunelli, S. A., Shair, H. N., & Hofer, M. A. (1994). Hypothermic vocalizations of rat pups (Rattus norvegicus) elicit and direct maternal search behavior. Journal of Comparative Psychology, 108(3), 298–303. https://doi.org/10.1037/0735-7036.108.3.298

Christensen, J., Grønborg, T. K., Sørensen, M. J., Schendel, D., Parner, E. T., Pedersen, L. H., & Vestergaard, M. (2013). Prenatal Valproate Exposure and Risk of Autism Spectrum Disorders and Childhood Autism. JAMA, 309(16), 1696. https://doi.org/10.1001/jama.2013.2270

Debiec, J., & Sullivan, R. M. (2017). The neurobiology of safety and threat learning in infancy. Neurobiology of Learning and Memory, 143, 49–58. https://doi.org/10.1016/j.nlm.2016.10.015

Deviterne, D., & Desor, D. (1990). Selective pup retrieving by mother rats: Sex and early development characteristics as discrimination factors. Developmental Psychobiology, 23(4), 361–368. https://doi.org/https://doi.org/10.1002/dev.420230407

Dufour-Rainfray, D., Vourc, P., & Tourlet, S. (2011). Fetal exposure to teratogens: Evidence of genes involved in autism. Neuroscience and Biobehavioral Reviews, 35(5), 1254–1265. https://doi.org/10.1016/j.neubiorev.2010.12.013

Ellenbroek, B. A., August, C., & Youn, J. (2016). Does prenatal valproate interact with a genetic reduction in the serotonin transporter? A rat study on anxiety and cognition. Frontiers in Neuroscience, 10(SEP), 1–10. https://doi.org/10.3389/fnins.2016.00424

Ferri, S. L., Abel, T., & Brodkin, E. S. (2018). Sex Differences in Autism Spectrum Disorder: a Review. Current Psychiatry Reports, 20(2), 9. https://doi.org/10.1007/s11920-018-0874-2

Green, J., Pickles, A., Pasco, G., Bedford, R., Wan, M. W., Elsabbagh, M., Slonims, V., Gliga, T., Jones, E., Cheung, C., Charman, T., Johnson, M., Baron-Cohen, S., Bolton, P., Davies, K., Liew, M., Fernandes, J., Gammer, I., Salomone, E., … McNally, J. (2017). Randomised trial of a parent-mediated intervention for infants at high risk for autism: longitudinal outcomes to age 3 years. Journal of Child Psychology and Psychiatry and Allied Disciplines, 58(12), 1330–1340. https://doi.org/10.1111/jcpp.12728

Gunnar, M. R. (2017). Social Buffering of Stress in Development: A Career Perspective. Perspectives on Psychological Science, 12(3), 355–373.https://doi.org/10.1177/1745691616680612

Hofer, M. A., & Shair, H. (1978). Ultrasonic vocalization during social interaction and isolation in 2-week-old rats. Developmental Psychobiology, 11(5), 495–504. https://doi.org/10.1002/dev.420110513

Hostinar, C. E., Johnson, A. E., & Gunnar, M. R. (2015). Early Social Deprivation and the Social Buffering of Cortisol Stress Responses in Late Childhood: An Experimental Study. Developmental Psychology, 51(11), 1597–1608. https://doi.org/10.1037/dev0000029

Kim, K. C., Lee, D. K., Go, H. S., Kim, P., Choi, C. S., Kim, J. W., Jeon, S. J., Song, M. R., & Shin, C. Y. (2014). Pax6-dependent cortical glutamatergic neuronal differentiation regulates autism-like behavior in prenatally valproic acid-exposed rat offspring. Molecular Neurobiology, 49(1), 512–528. https://doi.org/10.1007/s12035-013-8535-2

Li, Y., Zhou, Y., Peng, L., & Zhao, Y. (2017). Reduced protein expressions of cytomembrane GABA(A)Rβ3 at different postnatal developmental stages of rats exposed prenatally to valproic acid. Brain Research, 1671, 33–42. https://doi.org/10.1016/j.brainres.2017.06.018

Lord, C., Elsabbagh, M., Baird, G., & Veenstra-Vanderweele, J. (2018). Autism spectrum disorder. The Lancet, 392(10146), 508–520. https://doi.org/10.1016/S0140-6736(18)31129-2

Markram, K., Rinaldi, T., Mendola, D. La, Sandi, C., & Markram, H. (2008). Abnormal fear conditioning and amygdala processing in an animal model of autism. Neuropsychopharmacology, 33(4), 901–912. https://doi.org/10.1038/sj.npp.1301453

Mendez-Gallardo, V., & Robinson, S. R. (2014). Odor-induced crawling locomotion in the newborn rat: Effects of amniotic fluid and milk. Developmental Psychobiology, 56(3), 327–339. https://doi.org/10.1002/dev.21102

Moore, C. L., & Morelli, G. A. (1979). Mother rats interact differently with male and female offspring. Journal of Comparative and Physiological Psychology, 93(4), 677–684. https://doi.org/10.1037/h0077599

Moriceau, S., & Sullivan, R. M. (2006). Maternal presence serves as a switch between learning fear and attraction in infancy. Nature Neuroscience. https://doi.org/10.1038/nn1733

Nachmias, M., Gunnar, M., Mangelsdorf, S., Parritz, R. H., & Buss, K. (1996). Behavioral inhibition and stress reactivity: The moderating role of attachment security. CHILD DEVELOPMENT, 67(2), 508–522. https://doi.org/10.1111/j.1467-8624.1996.tb01748.x

Richmond, G., & Sachs, B. D. (1984). Maternal discrimination of pup sex in rats. Developmental Psychobiology, 17(1), 87–89. https://doi.org/ https://doi.org/10.1002/dev.420170108

Schneider, T., & Przewłocki, R. (2005). Behavioral alterations in rats prenatally to valproic acid: Animal model of autism. Neuropsychopharmacology, 30(1), 80–89. https://doi.org/10.1038/sj.npp.1300518

Sheldrick, R. C., Maye, M. P., & Carter, A. S. (2017). Age at First Identification of Autism Spectrum Disorder: An Analysis of Two US Surveys. Journal of the American Academy of Child and Adolescent Psychiatry, 56(4), 313–320. https://doi.org/10.1016/j.jaac.2017.01.012

Shionoya, K., Moriceau, S., Bradstock, P., & Sullivan, R. M. (2007). Maternal attenuation of hypothalamic paraventricular nucleus norepinephrine switches avoidance learning to preference learning in preweanling rat pups. Hormones and Behavior, 52(3), 391–400. https://doi.org/10.1016/j.yhbeh.2007.06.004

Stanton, M. E., & Levine, S. (1990). Inhibition of infant glucocorticoid stress response: Specific role of maternal cues. Developmental Psychobiology, 23(5), 411–426. https://doi.org/10.1002/dev.420230504

Stanton, M. E., Wallstrom, J., & Levine, S. (1987). Maternal Contact Inhibits Pituitary-Adrenal Stress Responses in Preweanling Rats. Developmental Psychobiology, 20(2), 131–145.

Suchecki, D., Rosenfeld, P., & Levine, S. (1993). Maternal regulation of the hypothalamic-pituitary-adrenal axis in the infant rat: the roles of feeding and stroking. Developmental Brain Research, 75(2), 185–192. https://doi.org/10.1016/0165-3806(93)90022-3

Wiedenmayer, C. P., Magarinos, A. M., McEwen, B. S., & Barr, G. A. (2003). Mother lowers glucocorticoid levels of preweaning rats after acute threat. Annals of the New York Academy of Sciences, 1008, 304–307. https://doi.org/10.1196/annals.1301.038

Zwaigenbaum, L., Bryson, S., Rogers, T., Roberts, W., Brian, J., & Szatmari, P. (2005). Behavioral manifestations of autism in the first year of life. International Journal of Developmental Neuroscience, 23(2-3 SPEC. ISS.), 143–152. https://doi.org/10.1016/j.ijdevneu.2004.05.001

